# Focused attention meditation changes the boundary and configuration of functional networks in the brain

**DOI:** 10.1101/664573

**Authors:** Shogo Kajimura, Naoki Masuda, Johnny King Lau, Kou Murayama

## Abstract

Research has shown that meditation not only improves our cognitive and motivational functioning (e.g., attention, mental health), it influences the way how our brain networks [e.g., default mode network (DMN), fronto-parietal network (FPN), and sensory-motor network (SMN)] function and operate. However, surprisingly little attention has been paid to the possibility that meditation alters the structure (composition) of these functional brain networks. Here, using a single-case experimental design with longitudinal intensive data, we examined the effect of mediation practice on intra-individual changes in the composition of whole-brain networks. The results showed that meditation (1) changed the community size (with a number of regions in the FPN being merged into the DMN after meditation), (2) changed the brain regions composing the SMN community without changing its size, and (3) led to instability in the community allegiance of the regions in the FPN. These results suggest that, in addition to altering specific functional connectivity, meditation leads to reconfiguration of whole-brain network structure. The reconfiguration of community structure in the brain provides fruitful information about the neural mechanisms of meditation.

## Introduction

Meditation is a practice aimed to enhance one’s core psychological capacities, such as attentional and emotional self-regulation (Tang, Hölzel, and Posner 2015). In several styles of practice, focused attention (FA) meditation involves sustaining attention to present-moment experiences without emotional reaction and judgment and has been found to produce significant beneficial outcomes, such as stress reduction (Goldin and Gross 2010) and improvements in attention processing (van den Hurk et al. 2010).

Past research indicates that FA meditation is primarily related to three brain networks: the fronto-parietal network (FPN), sensory-motor network (SMN), and default mode network (DMN) (Tang, Hölzel, and Posner 2015). The FPN mainly consists of the rostro- and dorso-lateral prefrontal cortex (lPFC), anterior insula, dorsal anterior cingulate cortex (dACC), and anterior inferior parietal lobule (aIPL); all of these brain areas are critical for cognitive control functions, such as regulation of attention and emotion (Dosenbach et al. 2008; Spreng et al. 2010; Sridharan, Levitin, and Menon 2008; Vincent et al. 2008). FA meditation, especially in the early stages of long-term practice, increases activation of FPN regions (Brefczynski-Lewis et al. 2007; Chiesa and Serretti 2010; Hasenkamp et al. 2012; Malinowski 2013), which is consistent with the general observation that focusing on the present moment requires effortful attentional control.

FA meditation also alters sensory experiences through the SMN (Berkovich-Ohana, Glicksohn, and Goldstein 2012; Cahn, Delorme, and Polich 2010), consisting of motor cortices, primary somatosensory cortex, and insula. In a previous study, these brain areas showed reduced activation in a four-day FA meditation when beginners meditated in the presence of noxious stimulation causing pain (Zeidan et al. 2011). This change in brain activity may be associated with enhanced body awareness, as FA meditation requires individuals to focus on a body part or internal experiences, such as breathing (Kabat-Zinn 2009).

The DMN, mainly consisting of the anterior medial prefrontal cortex (amPFC), posterior cingulate cortex (PCC), and posterior inferior parietal lobule (pIPL), is a network implicated in supporting spontaneous thoughts and self-referential processing (Christoff et al. 2016; Kajimura et al. 2018; Smallwood and Schooler 2015). Because sustained attention on an anchoring object (e.g., one’s breath) needs to detect distraction such as task-irrelevant thoughts, disengage attention from the distraction, and redirect attention on the object, the DMN is expected to be suppressed during FA meditation. In fact, the mPFC and PCC showed less activity during FA meditation, and functional connectivity between the PCC, dACC, and dlPFC was stronger in meditators compared to meditation-naive controls (Brewer et al. 2011). Increased functional connectivity between the PCC and task-positive regions was also observed in a different study (Grigg and Grady 2010). These results indicate that FA meditation may increase cognitive control over the DMN functioning (Brewer et al. 2011).

Although the previous work has provided various insights into how meditation influences the functional network of the brain, there are two critical limitations in the current literature. First, the brain networks were defined *a priori* in the previous studies, precluding the possibility that meditation practice can alter the structure of the primary brain networks themselves (i.e. FPN, SMN, and DMN). Because recent studies have shown that meditation can change functional connectivity across brain regions (Brewer et al. 2011; Hasenkamp et al. 2012; Kilpatrick et al. 2011; Taylor, Daneault, Grant, Scavone, Breton, Roffe-vidal, et al. 2013), the whole-brain composition of the FPN, SMN, and DMN may be altered as a consequence of meditation.

Second, most of the previous research has employed a one-shot pre-post or nonmeditator-meditator comparison design (Tang and Posner 2009; Zeidan et al. 2013), and compared the conditions after aggregating the data across heterogeneous participants. This inter-individual aggregation approach is useful to examine the effects of meditation averaged across participants. However, given the large individual differences in the whole-brain functional connectivity pattern (Bansal, Nakuci, and Muldoon 2018; Mueller et al. 2013), there is danger that the approach potentially masks important intra-individual changes in the composition of the brain networks (e.g., some participant-specific network structures may be canceled out by inter-individual aggregation). Therefore, adopting a design that allows us to focus on the intra-individual change may provide novel insights into how meditation alters the structure of the brain networks.

The current research aims to expand our understanding of meditation by addressing these two critical issues. For that purpose, we will examine the effects of FA meditation using a single-case experimental design with intensive longitudinal data. Single-case experimental designs have a long tradition in psychology (Fechner, Boring, and Winston 1966; Watson 1925), and in later years, they have been applied to intensive longitudinal data (for a systematic review, see Smith, 2013). Single-case experimental designs are effective in reliably detecting intra-individual changes in outcome variables in response to intervention (Smith 2013). However, this design has rarely been implemented in neuroimaging studies (for an exception without experimental manipulation, see Poldrack et al., 2015). Based on this design, we scanned a single participant repeatedly over a long period of time (65 days), employing FA meditation intermittently, and examined whether and how the whole-brain composition of the FPN, SMN, and DMN were altered on the days of FA meditation.

## Methods

### Participant

The participant (author S.K.) is a right-handed Asian male, aged 28 years and had no experience of meditation practice at the onset of the study. The participant is healthy with no history of neuropsychiatric disorders. The study was approved by the local institutional review board (UREC 16/28).

The participant underwent 65 scanning sessions, each on a different day between June 15th 2016 and November 11th 2016. The scanning time (between 9am – 5pm) was randomized across days. The data were not acquired on 6 out of the 65 days due to machine troubles and an additional 1 day was excluded due to excessive head movement (> 3.0mm). As a result, data from 58 days were used in the following analyses. On each day, the participant underwent a resting-state fMRI scan with eyes open for 10 minutes and completed two sets of questionnaires carried out for a different study. In 18 out of the 58 days (Fig. 1), the participant underwent a 15-min session of FA meditation practice a few minutes before scanning. In the practice, the participant were instructed to focus on his breathing, specifically on sensations of the breath on the nostrils, and to redirect attention from spontaneously upcoming thoughts to breathing when he realized his mind was wandered (Hasenkamp et al. 2012). Before the data collection, the participant studied the FA meditation several times from an auditory instruction developed by a professional trainer (Fujino et al., in revision) so that he did not need the instruction for each practice. In the following text, the “meditation condition” (MC) refers to the days on which scanning followed the FA meditation practice. The “no meditation condition” (NoMC) refers to the days on which there was no FA meditation practice prior to scanning.

**Figure 1.**
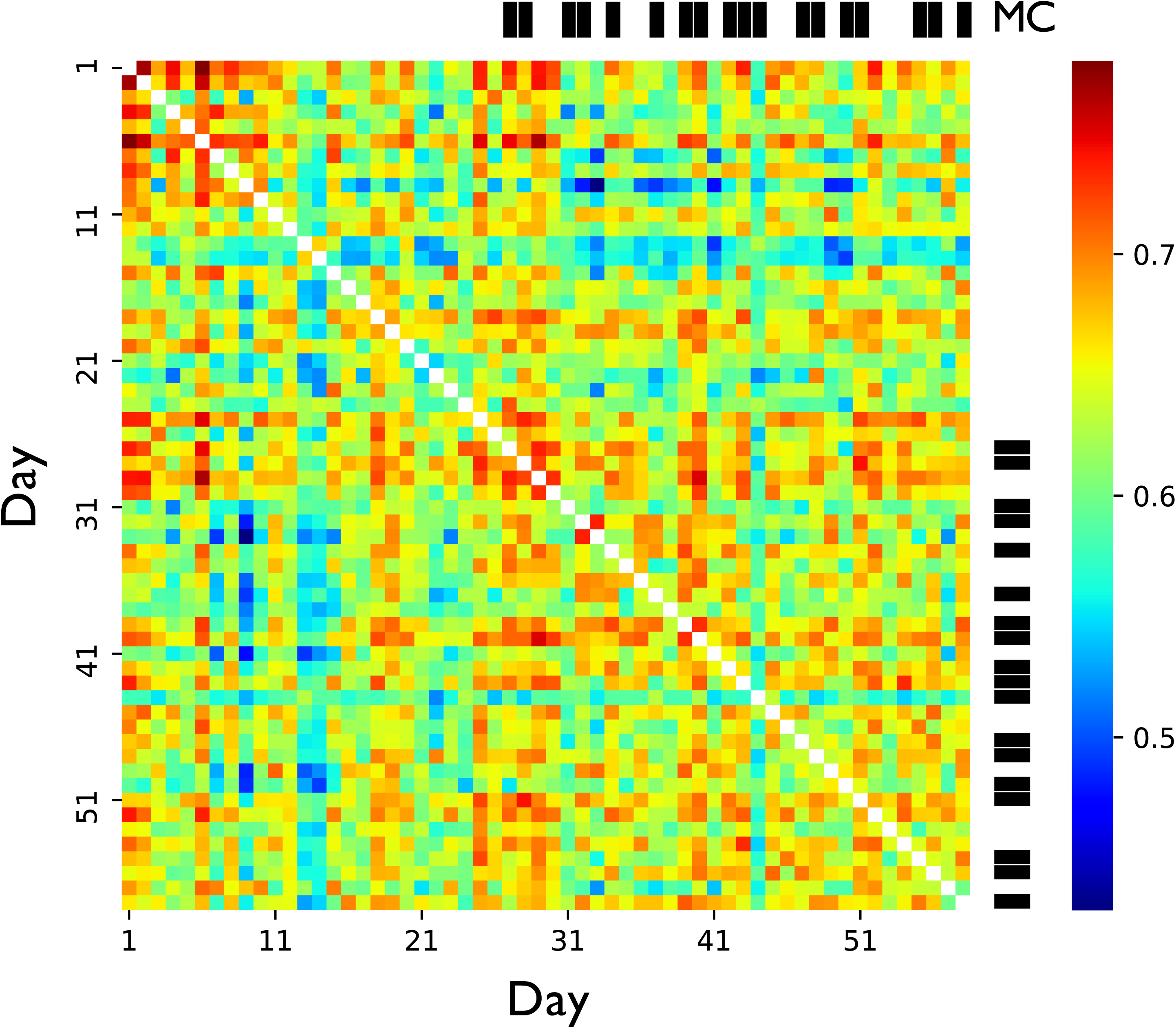
Similarity (correlation) between the functional networks for each pair of days. The color code represents the value of the correlation coefficient.

### Data acquisition

MR images were acquired using a Siemens 3.0-T Trio scanner equipped with a 32-channel head coil. The resting-state fMRI data were obtained using a single-shot, gradient-echo echo-planar imaging (EPI) sequence. Sequence parameters were as follows: repetition time/echo time (TR/TE) = 2,500/30 ms, slice thickness = 3.5 mm, field of view (FoV) = 256 mm, flip angle (FA) = 90°, data matrix = 64 × 64, in-plane resolution = 3.5 × 3.5 mm, 46 slices, 10 minutes scan length. Four dummy scans were discarded to remove the impact of magnetization instability. A high-resolution (spatial resolution: 1 × 1 × 1 mm) structural image was also acquired on the first day using a T1-weighted magnetization prepared rapid-acquisition gradient echo (MP-RAGE) pulse sequence.

### Data preprocessing

All preprocessing steps were performed using the Data Processing Assistant for Resting-State fMRI Advanced Edition (DPARSFA) (Chao-Gan and Yu-Feng 2010), which runs on Statistical Parametric Mapping 8 (SPM8) and the Resting-State fMRI Data Analysis Toolkit (REST) (Song et al. 2011). Data preprocessing included the following steps: realignment of all functional images using a six-parameter rigid body transformation; slice-timing correction to the middle slice of each volume; co-registration of the structural image (T1-weighted MPRAGE) to the mean functional image using a rigid-body transformation; segmentation of the transformed structural image into the gray matter, white matter, and cerebrospinal fluid (CSF); nuisance covariate regression of six head motion parameters, average white matter signals, CSF signals, and the global signal in native space; spatial normalization of the functional images to the Montreal Neurological Institute (MNI) stereotactic standard space; spatial smoothing of the functional images with a 6-mm full-width at half-maximum (FWHM) Gaussian kernel using the Diffeomorphic Anatomical Registration Through Exponentiated Lie Algebra (DARTEL) toolbox (Ashburner 2007); band-pass filtering (0.01–0.10 Hz) to reduce low-frequency drift and high-frequency physiological noise.

### Computation of functional connectivity

To calculate a functional connectivity matrix for each day, we defined the regions of interest (ROI) based on the Human Connectome Project (HCP)’s multi-modal parcellation version 1.0 (Glasser et al. 2016) and the automated anatomical labeling (AAL) atlas (Tzourio-Mazoyer et al. 2002). Of the 360 cerebral cortical ROIs defined by the HCP, one ROI (rh.R_8BL) was excluded because the obtained mask contained only 2 voxels. In addition to the remaining 359 cerebral cortical ROIs, we included 40 limbic and cerebellar ROIs defined by AAL, resulting in a total of 399 ROIs. For each ROI, we computed the spatial average of the signal within the mask. For each day, we quantified functional connectivity between each pair of the 399 ROIs by the absolute value of the Pearson correlation coefficient between the two fMRI time-course signals (Achard 2006; Fallani et al. 2014).

### Similarity of the functional connectivity across time

To calculate correlation of functional connectivity between days. Specifically, we first vectorized the functional connectivity between all pairs of 399 ROIs for each day into a 399 × 398 / 2 = 79,401 dimensional vector. Then, we computed the Pearson’s correlation coefficient between the two vectors for the corresponding days.

### Generalized Louvain method

The original Louvain method approximately maximizes the objective function (modularity) to partition the nodes in the given static network into communities. Communities are determined such that there are many edges within each community and relatively few edges between communities (Blondel et al. 2008). The generalized Louvain method considers the edges across multiple inter-dependent slices and optimizes the generalized modularity instead of separately optimizing the modularity for each slice. In the present study, a slice represents the static functional connectivity on one day. We ran the algorithm 100 times and selected the community structure yielding the largest generalized modularity value. As shown in the Results section, this procedure found four communities that are comparable with the communities in previous research (Meunier, Lambiotte, and Bullmore 2010; Moussa et al. 2012; Sporns 2013).

### Community labeling

First, we represented each community *i* (*i* = 1, 2, 3, 4) by its core members. The core members of the *i*th community were defined by the ROIs whose dominant community, that is the community to which the ROI belonged for the largest number of days under the given condition, was the *i*th community under both conditions. The relative overlap between the *i*th community and a community in the template communities defined by Yeo et al. (2011), *N_j_* (*j* = 1, 2, …, 7), was defined as 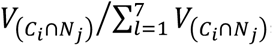, where *C_i_* is the set of voxels belonging to a core member of the *i*th community, *N_j_* is interpreted as the set of voxels belonging to mask *N_j_*, and *V_(C_i_∩N_j_)_* represents the number of voxels that belong to both *C_j_* and *N_j_*. We then labeled each community *i* according to the community in the template that exhibited the largest overlap with the *i*th community.

### Calculation and statistical significance test of community size

We defined the community size for a day *t* as the number of regions included in the time-dependent community on day *t* and tested their difference across condition by a 2 (practice: MC vs. NoMC) x 4 (community: Visual network, SMN, FPN, and DMN) analysis of variance (ANOVA).

### Calculation and statistical significance test of community coherence

We defined the similarity of each community *i* (*i* = 1, 2, 3, 4) between day *t*_1_ and day *t*_2_ by the Jaccard index, i.e. *J_(*X,Y*)_* = |X ∩ Y|/|X ∪ Y|, where *X* is the set of nodes in community *1* on day *t*_2_ and *Y* is the set of nodes in community *i* on day *t*_2_. Jaccard index *J_(X,Y)_* ranges between 0 and 1. One obtains *J_(X,Y)_* = 1 if and only if *X* and *Y* are exactly the same, and *J_(X,Y)_* = 0 if *X* and *Y* do not share any ROIs. For each of the four communities, we calculated the similarity between all pairs of 58 days, obtaining a 58 × 58 similarity matrix.

Given the similarity matrix for a community, we compared the coherence of the community within and across the practice conditions. Specifically, we adapted a permutation test, which is commonly used for testing the significance of single-subject research (Edgington 1980). The permutation test consists of the following three steps. In the first step, we classified the pairs of days into two groups. The congruent (CONG) group contained the pairs of days which both belonged to the MC or the pair of days which both belonged to the NoMC. In contrast, the incongruent (INCONG) group contained the pairs of days from the different conditions (i.e. one from the MC and the other from the NoMC). Because there were 18 MC days and 40 NoMC days, CONG and INCONG groups contained 933 and 720 pairs of days, respectively. In the second step, we computed the coherence of the community, which is a Welch’s *t*-value, by comparing the averaged similarity value between the CONG and INCONG groups. We then randomized the days by reassigning 18 uniformly randomly selected days to the fictive MC and the remaining 40 days to the fictive NoMC, and calculated the coherence for the randomized data. We repeated this procedure 10,000 times to obtain the null distribution of the coherence for the randomized labeling. In the third step, we assessed the probability of obtaining the coherence calculated on the basis of the true labeling of the days (i.e., MC or NoMC) or more extreme coherence under the null model in which the MM condition was randomly assigned to individual days.

### Calculation and statistical significance test of flexibility

We defined the flexibility of a ROI under each condition using the inverse participation ratio (IPR) (Derrida and Flyvbjerg 1987). The IPR of a ROI under a condition is defined as

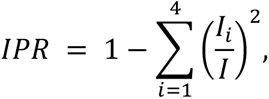

where *I* represents the number of days (NoMC, 40; MC, 18), and *I_i_* represents the number of days in which the ROI belonged to community *i* (*i* = 1, 2, 3, 4). The IPR is equal to 0.75, which is the largest possible value, when a ROI belongs to all the communities with the same probability. In this case, the ROI is the most flexible in terms of the community membership. The IPR is equal to 0, which is the smallest possible value, when the ROI belongs to the same single community in all the days. In this case, the ROI is the least flexible. To investigate whether the FA meditation affects the community-wide flexibility of ROIs, for each community, we applied a paired sample *t*-test to test the mean difference in the flexibility between the two conditions. In this particular analysis, we defined each community *i* by its core members, i.e., the ROIs that belonged to community *i* as the dominant community under both conditions.

## Results

### Similarity of the functional connectivity across time

To examine variability of the functional connectivity over time, we calculated correlation of functional connectivity between days (Fig. 1). The correlation value ranged from 0.429 to 0.780, suggesting that the functional connectivity of a single person varies on a daily basis. This result is consistent with previous longitudinal scanning data from a single participant (Poldrack et al. 2015).

### Finding community structure

To detect the community structure in the time-varying functional connectivity across the 58 days, we applied a generalized Louvain method (Jeub et al. 2017) (Fig. 2). To label the four communities detected in the current study, we assessed how these communities overlapped with the template communities defined by Yeo et al. (2011). As the result, we labeled the four communities as visual network (80.8% overlap), SMN (50.3% overlap), FPN (45.3% overlap), and DMN (58.4% overlap) (diagonal in Fig. 3A).

**Figure 2.**
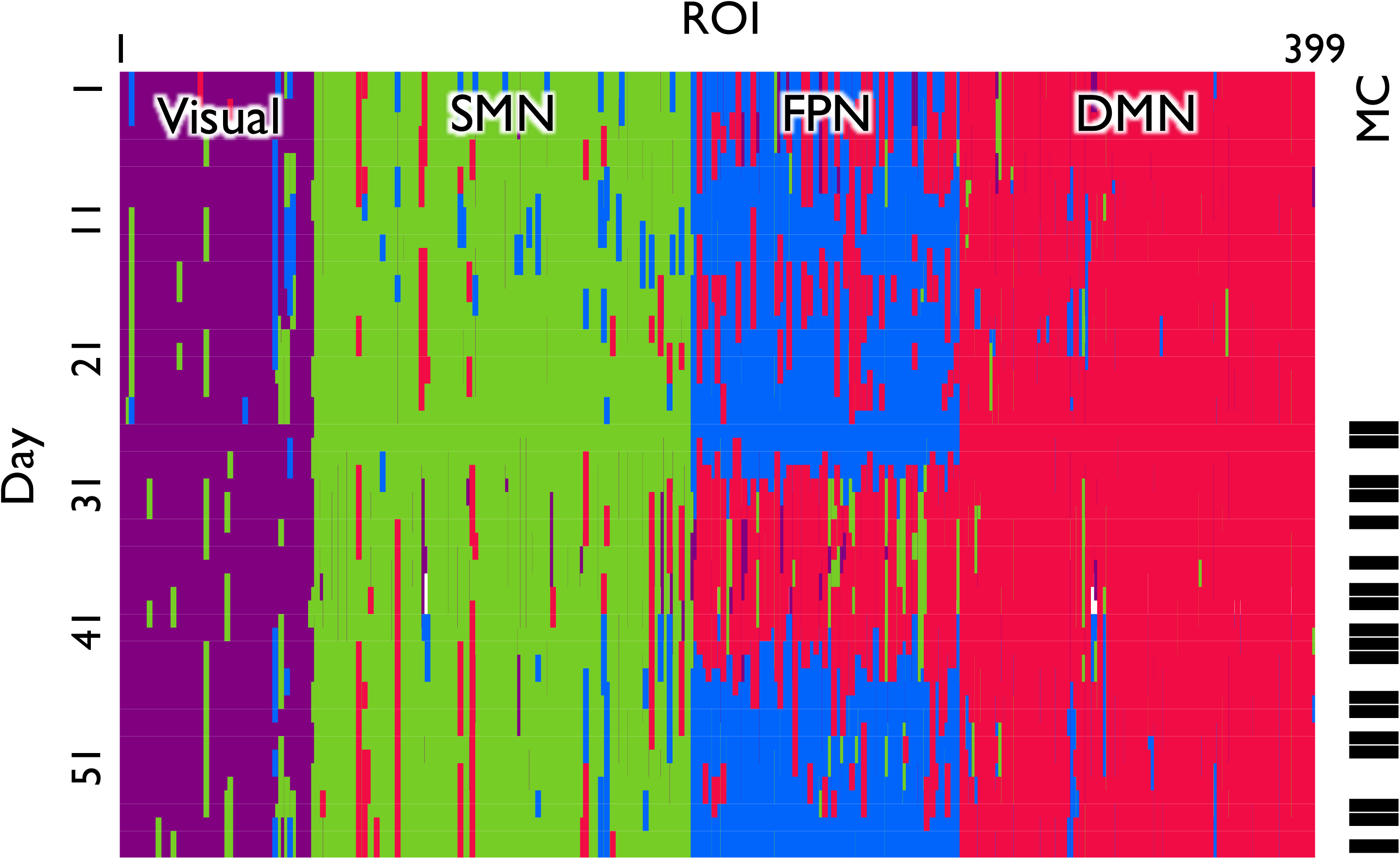
Time-dependent community structure. The rows and columns correspond to the days and ROIs, respectively. The meditation days are colored in black in the rightmost column. The four communities were labeled the visual network (colored in purple), sensory-motor network (SMN; green), fronto-parietal network (FPN; blue), and default mode network (DMN; red). A white stripe describes a day in which a ROI belonged to more than one community and could not be assigned a community label.

**Figure 3.**
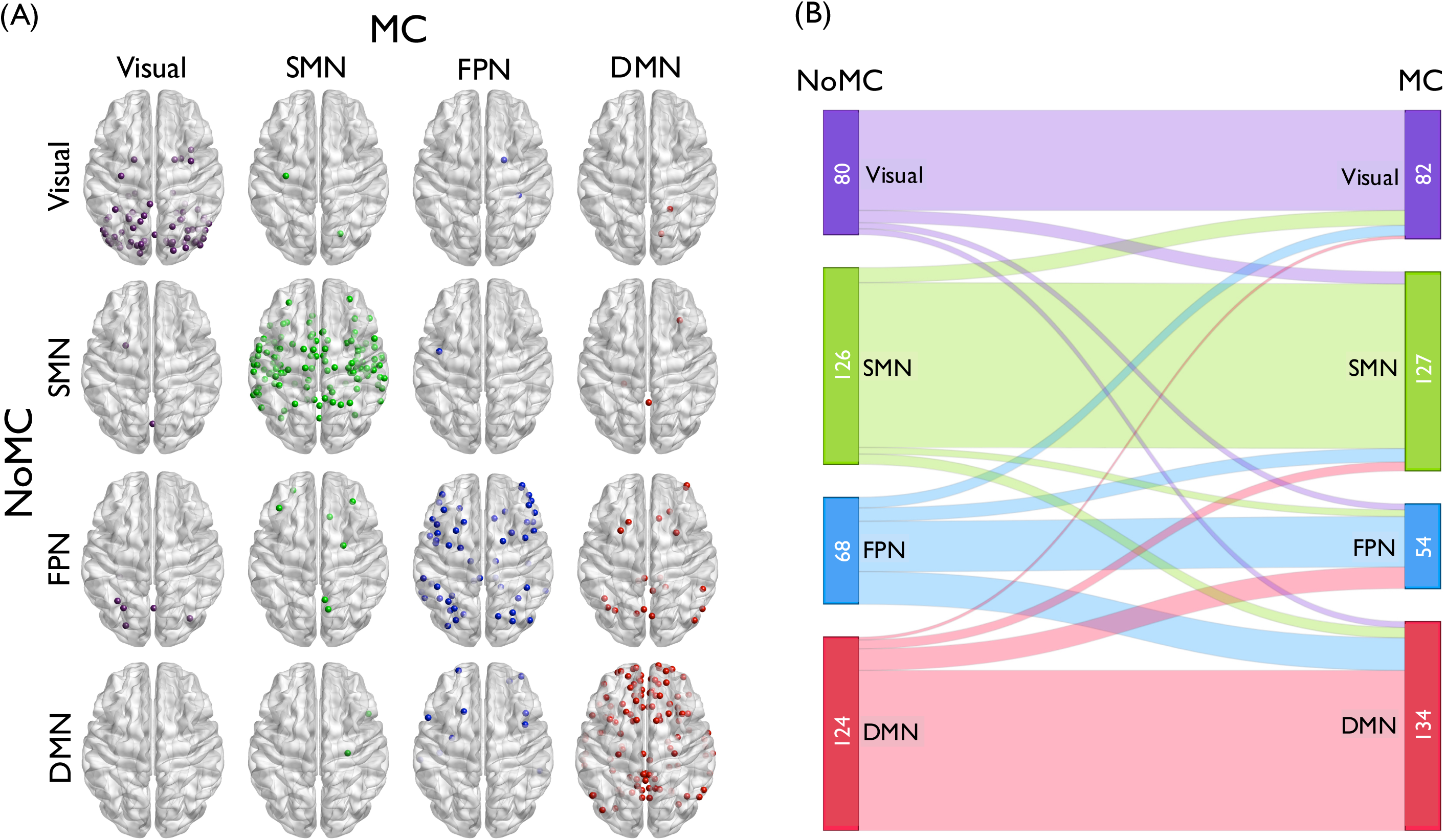
The change of dominant community affiliation between the NoMC and MC. (A) The shift of the dominant community allegiance of ROIs between the conditions. The schematic pictures of the brain on the diagonal show the ROIs that belonged to the same community with the highest probability across the two conditions. Those off the diagonal show the ROIs that belonged to different communities in the two conditions. (B) Quantitative description of dominant community shift across the conditions.

### Metrics that quantify the changes in community structure

We examined the effect of FA meditation practice on intra-individual changes in the composition of the whole-brain networks with three metrics: *community size*, *community coherence*, and *flexibility*.

### Community size

The ANOVA showed a significant main effect of community (*F*_(3, 224)_ = 133.2, *p* < 0.001) while no significant main effect was observed for the condition (*F*_(1, 224)_ = 0.01, *p* = 0.911). The interaction between the community and the condition was statistically significant (*F*_(3, 224)_ = 2.73, *p* = 0.044), suggesting that there was a significant change in the relative community size between conditions. A simple main effect analysis showed that the FPN community tended to be smaller (*F*(1, 56) = 2.317, *p* = 0.134) and the DMN community tended to be larger (*F*_(1, 56)_ = 2.515, *p* = 0.118) under the MC compared with NoMC.

Fig. 3A shows whether the ROIs stayed in the same community or changed to a different dominant community between the two conditions (i.e. represents the main community that a ROI belonged to under each condition). The on-diagonal brains show the ROIs that stayed in the same dominant community across the two conditions. The off-diagonal brains show the ROIs that belonged to different dominant communities between the two conditions. Consistent with the ANOVA results, the figure shows that a large number of ROIs (i.e. 21 ROIs) that belonged to the FPN in the NoMC shifted to the DMN in the MC (row 3, column 4; other community pairs ≤ 11 ROIs). The shift in the dominant community affiliation between the conditions is quantitatively depicted in Fig. 3B.

### Coherence of community composition

The community size is one way of examining the change in the composition of the brain network across the two conditions. In fact, even if the relative community size is the same between the two conditions, the constituent ROIs of each community may be substantially different in the two conditions. Therefore, for each community, we examined the extent to which the community as identified by the set of ROIs comprising it is stable within each of the two conditions (community coherence). The results of the permutation test are shown in Fig. 4. For the SMN, FPN, and DMN, the coherence of the community structure within the same condition (CONG) was larger than the coherence between the NoMC and MC (INCONG; SMN, *p* = 0.030; FPN, *p* = 0.028; DMN, *p* = 0.007). This result indicates that FA meditation practice has changed the community composition (i.e., the set of ROIs composing the community) in the SMN, FPN, and DMN. The results for the FPN and DMN are a direct consequence of our findings on the community size because a change in the community size implies a decrease in the coherence value. In contrast, the composition of the SMN also changed despite its stable size between the two conditions. For the visual network community, there was no difference in the coherence (*p* = 0.137), which implies that the set of ROIs composing the visual network community was not significantly influenced by FA meditation.

**Figure 4.**
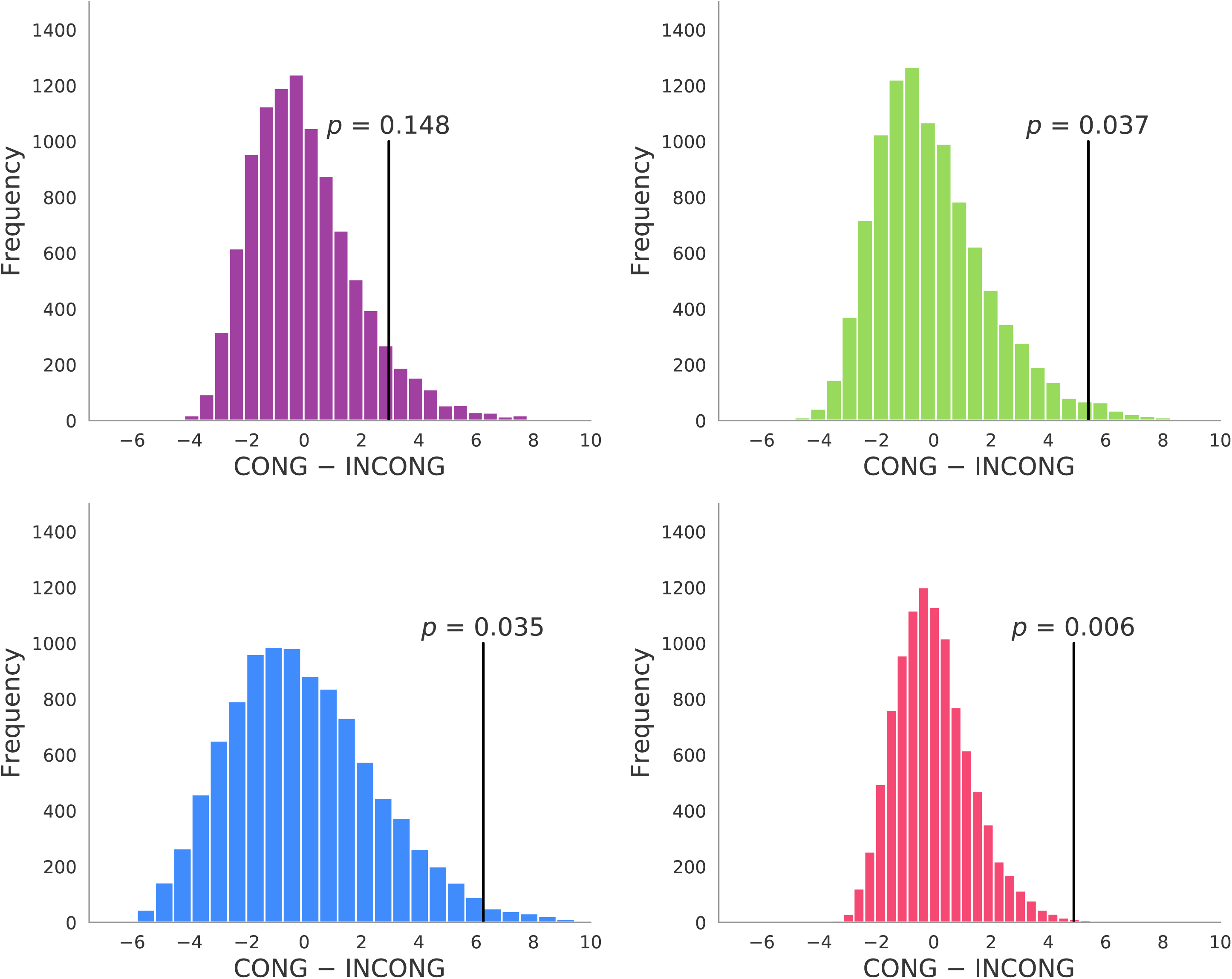
Coherence of the community within and across conditions. Coherence is a Welch’s *t*-value that represents the difference in averaged similarity value of a community structure between two groups, i.e., the congruent (CONG) and incongruent (INCONG) group. The CONG group contained pairs of days that both belonged to the MC or the NoMC. The INCONG group contained the pairs of days from the different conditions (i.e. one from the MC and the other from the NoMC). The figure shows null distributions of coherence (distributions of coherence for permutated data); vertical lines represent the *t*-values observed with the true labeling and their corresponding *p-*values representing the probability of obtaining such observed *t*-values (or more extreme *t*-values) in the null distribution.

### Flexibility of community allegiance

To assess experience-related changes in community allegiance of a ROI, we defined the flexibility of a ROI under each condition using the inverse participation ratio (IPR) (Derrida & Flyvbjerg, 1987). The change in flexibility between MC and NoMC for individual ROIs belonging to each community (i.e. Visual network, SMN, FPN, and DMN) is shown in Fig. 5. Positive values mean that flexibility of the ROI increased as a consequence of FA meditation. One-sample *t*-tests of the difference in flexibility between the two conditions for each community revealed that FA meditation significantly enhanced flexibility of the ROIs in the FPN (M = 0.17, SD = 0.13, *t* = 7.334, *p* < 0.001). Other communities did not show a significant difference in flexibility of the ROIs (|M| ≦ 0.02, SD ≧ 0.10, |*t*| ≦ 0.802). These results suggest that FA meditation increases the flexibility of the FPN community, but not the visual network, SMN, or DMN. Table 1 summarizes the results.

**Figure 5.**
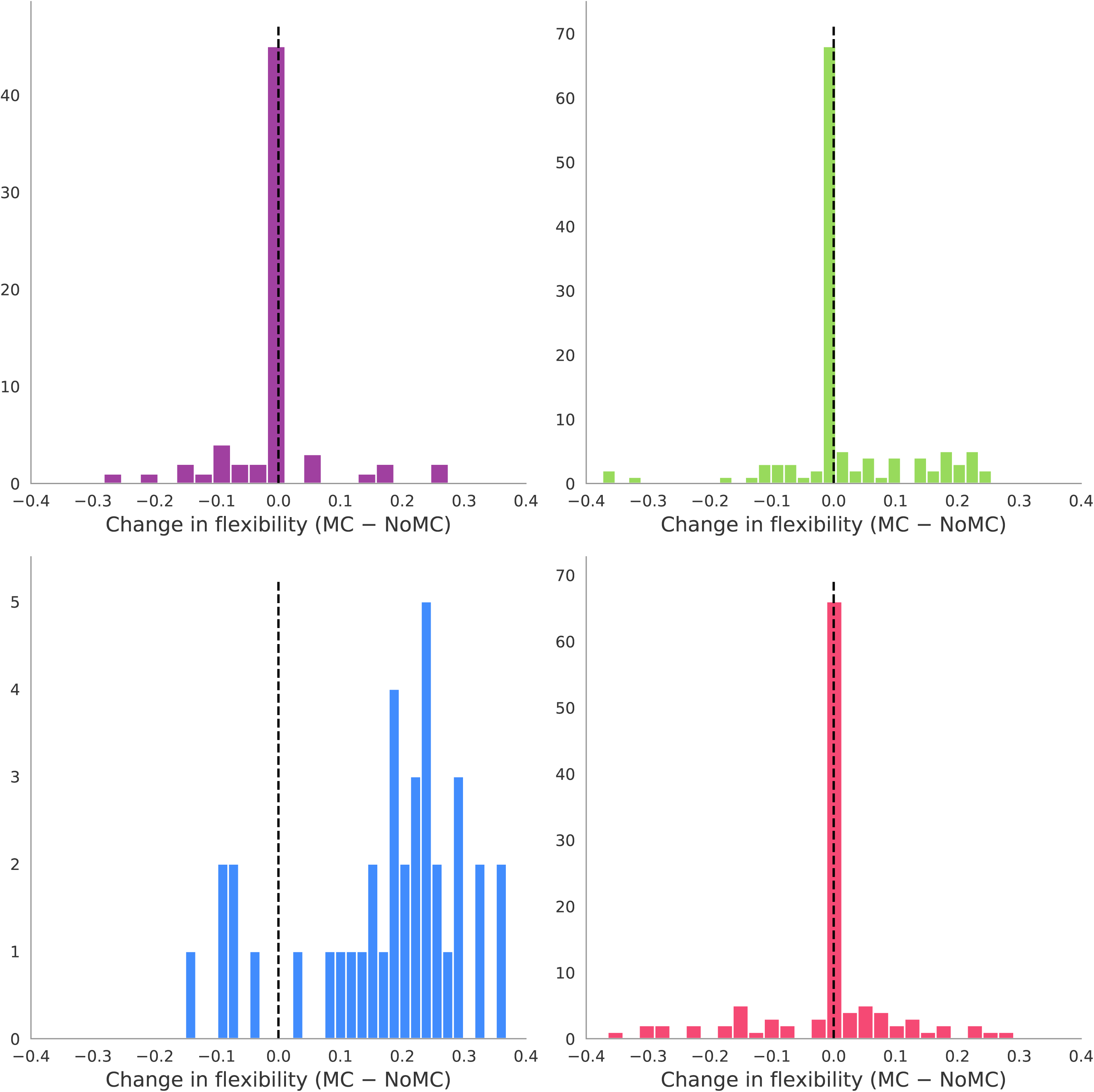
Changes in flexibility of ROIs between two conditions. Histograms for each community show the distribution of the values of ROIs that belonged to the corresponding community in both conditions.

**Table 1.** Summary of the meditation effect. *n.s.* indicates not significant.

| Network | Size | Coherence | Flexibility |
| --- | --- | --- | --- |
|  | The number of ROIs in the community | Change in the composition of ROIs in a community | Frequency with which ROIs change their community allegiance |
| Visual | *n.s.* | *n.s.* | *n.s.* |
| SMN | *n.s.* | NoMC ≠ MC | *n.s.* |
| FPN | NoMC > MC | NoMC ≠ MC | NoMC < MC |
| DMN | NoMC < MC | NoMC ≠ MC | *n.s.* |

## Discussion

Previous studies have provided accumulating evidence that FA meditation changes activation and connectivity patterns between specific brain regions (Brewer et al. 2011; Grigg and Grady 2010; Hasenkamp and Barsalou 2012; Kilpatrick et al. 2011; Tang and Posner 2009; Taylor, Daneault, Grant, Scavone, Breton, Roffe-Vidal, et al. 2013; Zeidan et al. 2013). Extending previous work, we employed a whole-brain graph theoretic analysis with a single-case experimental design using intensive longitudinal data to reveal that FA meditation provokes the reconfiguration of the community structure of the whole-brain functional network.

We found that the size of the FPN decreased and that of the DMN increased as a consequence of FA meditation. This result is consistent with the previously shown enhanced connectivity between the DMN and the FPN (Brewer et al. 2011; Grigg and Grady 2010). Brewer and colleagues (2011) reported that experienced meditators showed increased functional connectivity of the PCC with the dACC and dlPFC both during rest and meditation. Similarly, Grigg and Grady (2010) reported increased functional connectivity between the PCC and task-positive brain regions. Based on these findings, Brewer et al. (2011) proposed that the composition of these brain networks might have changed over time and become a new “default mode” that can be observed during meditation as well as during the resting state (Brewer et al., 2011). The current observation that some ROIs shift from the FPN to the DMN after FA meditation supports their hypothesis.

The FPN, which is one of the critical networks related to FA meditation (Tang, Hölzel, and Posner 2015), showed enhanced flexibility under the FA meditation condition. A previous study suggested that enhanced flexibility in the FPN may reflect an (initial) learning process of a task (Bassett et al. 2011). Accordingly, although the previous study used a different task with a different time scale, i.e., Bassett et al. (2011) observed the change in the flexibility within a few hours of motor-task training, the increased flexibility in the FPN in the current study may indicate that some form of learning process is operative in FA meditation (e.g., how to control breathing, attention, bodily sensations etc.). Future studies should directly examine whether the enhanced flexibility in the FPN is related to a long-term learning process in FA meditation. If this hypothesis is confirmed, the flexibility of community allegiance in the FPN may be used as a biomarker to assess the efficacy and the progress of FA meditation.

Although the SMN showed a change in coherence across the two conditions, as the FPN and DMN did, the size and flexibility of the SMN were intact. In this sense, FA meditation influenced the SMN but to a lesser extent than it influenced the FPN and DMN. This weaker alteration in the SMN might be because awareness of the body sensation that recruits the SMN (Zeidan et al. 2011) is not a primary target of FA meditation in contrast to the attentional enhancement, which is associated with the FPN, and reduction of mind wandering, which is associated with the DMN. One can test this possibility in future research by using an external stimulus (e.g., a fixation) as an object on which attention should be focused during FA meditation rather than a body sensation.

The present study demonstrated the value of a single-case experimental design with intensive longitudinal data. It allows us to detect intra-individual changes in the whole-brain network composition without being influenced by the large heterogeneity of individuals’ brain functional networks (Mueller et al. 2013). Previous studies on meditation heavily relied on the expert-beginner and/or pre-post comparison design (Tang, Hölzel, and Posner 2015). Future research should be encouraged to adopt the single-case research design more frequently to seek further insights into intra-individual changes in patterns of brain networks as a consequence of meditation. One obvious limitation of the current research design is that the data were collected from a single participant, which makes it impossible to examine potential individual differences in our findings. However, although research in cognitive neuroscience typically collects data from multiple participants, for the majority of studies, their main focus is on the aggregated pattern of the brain activation/connectivity (but see person-centered research, e.g., Bansal, Nakuci, and Muldoon, 2018), and individual differences have been typically treated as random noise (sampling error). Therefore, in our view, this limitation is superseded by the strength of the current design: sensitivity to the nuanced intra-individual changes in brain signals and functional connectivity. Nevertheless, the potential of the current intensive longitudinal design would be considerably improved by data obtained from multiple participants in future studies. Another limitation of the present study is that the participant performed meditation practice only for three months, which is considerably shorter than the previous studies with experts (e.g., more than one year of regular practice) (Hasenkamp et al. 2012). Future research should collect data for a more prolonged period of time to examine how the progress of practice induces long-term changes in the community structure.

## Acknowledgements

This research was supported by the Marie Curie Career Integration Grant, Award Number CIG630680; JSPS KAKENHI (Grant Numbers 15H05401; 16H06406, 18H01102; 18K18696), F. J. McGuigan Early Career Investigator Prize from American Psychological Foundation; and the Leverhulme Trust (Grant Numbers RPG-2016-146 and RL-2016-030).

## Data Availability Statement

The data that support the findings of this study are available from the corresponding author upon reasonable request.

